# In-situ calibration of passive samplers for monitoring host-associated fecal markers in an urban river

**DOI:** 10.1101/2025.06.30.662257

**Authors:** Yawen Liu, Rory Verhagen, Metasebia Gebrewold, Wendy J.M. Smith, Darryl W. Hawker, Stuart L. Simpson, Warish Ahmed

## Abstract

Passive sampling in aquatic environments has shown promise as a time- and cost-efficient method with improved sensitivity for detecting microbial pollution in dynamic and variable conditions. However, quantitative descriptions of its field performance have been scarce, impeding its broader application in environmental settings. The performance of two membrane-based passive samplers (Torpedo and MSTFlow) in quantifying human (*Carjivirus*, Pepper Mild Mottle Virus [PMMoV], Tomato Brown Rugose Fruit Virus [ToBRFV]) and avian (*Helicobacter* spp. GFD marker) fecal pollution markers in an urban riverine environment over a 72-h deployment was evaluated using a first-order kinetic model to characterize their microbial adsorption characteristics. Results were compared with those from parallel composite and time-weighted auto-sampling. Accumulation in passive samplers exhibited an initial lag phase (0 to 8 h for viruses) and reached equilibrium within 16 h of initial deployment, except for GFD. Sampling rates were highest for PMMoV (6.29 and 5.11 mL/h) for MSTFlow and Torpedo samplers respectively, followed by *Carjivirus* (4.63 and 3.63 mL/h), ToBRFV (2.17 and 1.22 mL/h) and GFD (0.68 mL/h in Torpedo sampler). At equilibrium, both sampler configurations accumulated more than 22-170 times the amount of target gene copies in each mL of grab water samples depending on sampler and virus. These findings highlight passive sampling as a promising, sensitive, cost-effective, and low-maintenance alternative for continuous microbial water quality monitoring in aquatic environments. Overall, this study advances the understanding of passive sampling kinetics and supports broader adoption of passive samplers for tracking fecal pollution in environmental waters.

## 1. Introduction

Traditional water quality methods to detect/quantify fecal indicators and pathogens in aquatic environments have long relied on discrete grab sampling and large-volume filtration approaches (Dorevitch et al., 2011; Hozalski et al., 2024). While these have been useful for pathogen monitoring, grab samples provide only a brief snapshot of microbial water quality that may not be representative of characteristics over longer time frames, and may fail to detect pathogens present in low concentrations (Rafiee et al., 2021; Bivins et al., 2022). Additionally, grab samples can be prohibitively expensive if repeated sampling and analytical effort is required. As such, there is a growing need for alternative sampling approaches that can overcome these drawbacks while accommodating the dynamic nature of microbial contamination in environmental waters. Passive sampling offers a promising alternative, as a sample-cost and labour time-efficient technique, that may enable time-integrated monitoring of fecal indicators and pathogens (Karamati et al., 2024).

Passive sampling refers to the process of capturing contaminants from water over extended periods without the need for energy input, such as provided by pumps or mechanical systems (Górecki and Namieśnik, 2002; Salim and Górecki, 2019). Passive sampling methods typically use adsorbent materials that collect contaminants, such as pathogens, making them particularly useful for providing time-integrated monitoring in aquatic environments where conventional grab sampling or use of expensive autosamplers is impractical (Karamati et al., 2024). Passive sampling for microorganisms dates back to the mid-20th century. Pioneering work by Moore in 1948 introduced the “Moore swab,” a simple gauze swab attached to a string, which became a practical method for detecting pathogens in water (Moore, 1951).

Despite this early development, progress in passive sampling for microbial monitoring was limited until 2020, when the Coronavirus Disease 2019 (COVID-19) pandemic reignited interest in its application for wastewater surveillance. At this time, the technique demonstrated its potential for early detection of Severe Acute Respiratory Syndrome Coronavirus 2 (SARS-CoV-2) in sewage catchments (Corchis-Scott et al., 2021; Schang et al., 2021; Habtewold et al., 2022). For SARS-CoV-2 RNA, the detection frequency and concentration on passive samplers were largely influenced by viral loads in wastewater and sampler deployment duration (Schang et al., 2021; Hayes et al., 2024). Passive sampling using materials such as negatively charged membranes, Moore swabs and cheesecloth has shown similar or superior detection frequency of SARS-CoV-2 RNA compared to grab or composite sampling (Hayes et al., 2021; Rafiee et al., 2021; Schang et al., 2021). Negatively charged membranes have generally performed well for passive sampling in terms of reproducibility and sensitivity and been found to be effective for both detection and quantification of viruses in wastewater (Hayes et al., 2021; Schang et al., 2021; Bivins et al., 2022; Karamati et al., 2024).

Passive sampling is intended to provide a cumulative measure of contaminants over time, reducing any temporal and spatial fluctuations observed from analysis of discrete grab samples and improving the detection limits of pathogens with very low abundance (Payment and Locas, 2010; Karamati et al., 2024). A recent study compared a torpedo-shaped 3D printed passive sampler with composite sampling for detecting *Escherichia coli* (*E. coli*) and *Cryptosporidium* spp. in wastewater and surface water (Law et al., 2025). Both sampling methods detected *E. coli* at all time points (4 to 96 h). The detection frequencies of *Cryptosporidium* spp. were comparable between passive and composite sampling in surface water (31% vs. 41% of samples) and wastewater (76% vs. 86%). In this work, *E. coli* showed linear accumulation on passive samplers up to 96 h in surface water and 24 h in wastewater while *Cryptosporidium* spp. accumulation was linear up to 96 h in both matrices. These findings suggest that passive sampling is an emerging method that complements traditional active approaches for pathogen monitoring in aquatic environments.

While passive sampling holds promise for monitoring fecal indicators and pathogens in environmental waters, accurate quantification remains a significant challenge. Different passive sampling materials immersed in a particular water column result in different microbial (or their DNA/RNA signatures) accumulation rates over time (Corchis-Scott et al., 2021). A deeper understanding of the sampling dynamics involved in passive samplers is essential to exploit their effective application in monitoring microbial analytes in aquatic systems (Hayes et al., 2024).

The primary objectives of this study were to: (i) evaluate the performance of two membrane-based passive samplers (Torpedo and MSTFlow) in detecting human (*Carjivirus*, pepper mild mottle virus (PMMoV), tomato brown rugose fruit virus (ToBRFV), and avian (*Helicobacter* spp. associated (GFD)) fecal markers in an urban riverine environment at various time points within a 72 h deployment window; (ii) analyze the marker accumulation kinetics of the passive samplers using a first-order kinetic model to estimate the sampling rate and equilibrium time; and (iii) compare the passive sampling method to traditional sampling approaches (grab, time-weighted and composite) in terms of host-associated fecal marker detection and quantification. Torpedo samplers have been previously applied in riverine and wastewater environments and have demonstrated utility in capturing microbial targets. In contrast, MSTFlow is a newly developed sampler with a design similar to the Torpedo, featuring a modified casing. This study aims to advance the application of passive sampling techniques in environmental water quality surveillance, contributing to the development of more efficient, cost-effective, and continuous monitoring solutions for microbes in aquatic environments.

## 2. Materials and methods

### 2.1. Passive samplers

Two passive samplers were used in the study: (i) Torpedo, and (ii) MSTFlow. The Torpedo-style sampler is reusable and has been widely utilized for wastewater surveillance of SARS-CoV-2 and other viruses and has been demonstrated to achieve efficient capture of viral particles over time in flowing and static water systems (Schang et al., 2021). The housing material of this sampler was constructed from plastic, 3D-printed in the shape of a torpedo (cylindrical with a streamlined pointed front end) to help it remain stable in flowing water and to minimize the uptake of larger solid particles > 5mm (Law et al., 2025). The housing has a network of small orifices that allow wastewater to enter and interact with the adsorption materials such as membranes, gauze and swabs housed inside (Supplementary Figure SF1a).

The MSTFlow sampler was originally designed to capture viruses from underfloor wastewater effluent drains and to house up to three membrane-based sorbents separated within internal chambers. The sampler housing is cylindrical with cones at both ends, and also comprises a 3D-printed plastic material (liquid crystal polymer) containing a network of orifices (Supplementary Figure SF1a). The diameter of the MSTFlow sampler is wider than that of the Torpedo sampler and the orifices larger to facilitate a less restricted flow of liquids through the structure. The design was specifically intended to enable water to interact with the separate membranes on all sides when the sampler is placed into a drain. The membranes inside the sampler are held separated (so as not to overlap) by an insert which secures the edge of the membranes to the housing. For the Torpedo design, membranes were frequently observed to be overlapping at time of collection owing to movement following deployment. The MSTFlow design also enabled easier collection of the membranes by opening the housing using a simple twist-lock at either end of the cones, a feature that simplified the assembly and disassembly procedure.

A negatively charged membrane (0.45-μm mixed cellulose ester membrane; Millipore, REF#HAWP04700, Darmstadt, Germany) was chosen as the passive sampling material for both types of samplers. These membranes have been shown to be capable of adsorbing microorganisms from water facilitating nucleic acid extraction and quantification using molecular methods such as quantitative polymerase chain reaction (qPCR) or reverse transcription qPCR (RT-qPCR) while avoiding the accumulation of inhibitors and have yielded promising results for wastewater surveillance of pathogens (Vincent-Hubert et al., 2017; Habtewold et al., 2022; Hayes et al., 2024).

### 2.2 Passive sampler deployment and sample collection

Passive samplers (Torpedo and MSTFlow; each with two membranes) were deployed in an urban river on December 4, 2024, downstream of a sewage infrastructure pipeline that transports untreated wastewater from large populations (∼150,000 people) across a large tidal river to a resource recovery centre for treatment. The precipitation data from November to December 2024 in the study area were taken from the Bureau of Meteorology (BOM) site (http://www.bom.gov.au/, accessed on March 12, 2025) and provided in Supplementary Table ST1. Water quality at the deployment site is summarised in Table ST2.

Both passive samplers were deployed in triplicate for durations of 1, 4, 8, 12, 24, 48 and 72 h. All samplers were attached to a large metal mesh structure with pairs of Torpedo and MSTFlow positioned in close proximity to enable performance comparison. Care was taken to ensure that their placement did not interfere with water flow over their respective surfaces. The mesh securing the 42 samplers was submerged horizontally 0.5 m below the river water surface. At each time point following deployment, three pairs of samplers were removed for analysis (Supplementary Figure SF1b).

During the deployment period, complementary grab samples were also taken from the river on initial deployment of the samplers and thereafter at the same time points (i.e., 1, 4, 8, 12, 24, 48 and 72-h). The samples were collected in 500 mL sterile plastic containers using a telescopic sampling device to submerge and fill the bottle at 0.5 m depth. Aliquots of 100 mL from each grab sample were pooled to create a 72-h composite water sample. An autosampler constructed in-house (Automated Aquatic Sample Sipper, referred to hereafter as ‘Sipper’) was used to collect 24-h time-weighted river water samples (sampling every 15 s, measuring 85 to 98 mL in total) over a 24-h deployment period. Three consecutive samples were intended to be collected, representing the intervals: 0 to 24 h, 24 to 48 h, and 48 to 72 h. In practice, the final sample was not collected due to a malfunction in the Sipper.

All samples were transported on ice to the laboratory and stored in a refrigerator at 4 °C on the day of collection. One membrane from each passive sampler was carefully removed using sterile tweezers, rolled, transferred into 7 mL bead-beating tubes (Qiagen, Hilden, Germany), and stored at -20°C. The remaining membrane was archived in a sterile container at -20°C. Grab, 24 h time-weighted and composite water samples were stored at 4°C. All samples were analyzed within 72 h of collection.

### 2.3. Sample concentration and nucleic acid extraction

A modified adsorption-extraction (AE) concentration method was employed for the concentration of bacteria and viruses from grab, composite, and 24 h time-weighted water samples (Akter et al., 2024). A total of 300 mL from individual grab and composite water samples, and the two 24 h time-weighted samples (aliquoted at 98 mL and 85 mL) were filtered through MF-Millipore 0.45-µm MCE membranes (47 mm; Cat no. HAWP04700) (Millipore, Burlington, Massachusetts, USA) via a filter flask (Merck Millipore Ltd.). After filtration, the membrane was rolled and transferred into a 7 mL bead-beating tube (Qiagen) and stored at -20°C alongside the membranes from passive samplers.

A Qiagen RNeasy PowerWater Kit (Cat. No. 14700-50-NF, Qiagen) was used for nucleic acid extraction directly from the intact membranes. The DNase treatment step was omitted from the extraction protocol to obtain total nucleic acid from each sample. The nucleic acid samples were then eluted in 150 μL of nuclease-free water and stored at -20°C prior to qPCR and RT-qPCR analysis.

### 2.4. qPCR and RT-qPCR assays

Previously published qPCR and RT-qPCR assays were used for the analysis of host-associated fecal markers, *Carjivirus* (Stachler et al., 2017), PMMoV (Rosario et al., 2009; Haramoto et al., 2013), ToBRFV (Natarajan et al., 2023), and the GFD marker (Green et al., 2012). The sequences of primers and probes as well as cycling parameters for all assays are provided in Supplementary Table ST3.

qPCR amplifications were undertaken in 20 μL reaction mixtures using 10 μL of 2X QuantiNova Probe PCR Master Mix (Qiagen) and 3 μL of template. RT-qPCR amplifications were performed in 20 μL reaction mixtures using 5 μL of TaqMan Fast Virus 1-Step Master Mix (Applied Biosystem, California, USA) and 2 μL of template. Additionally, the qPCR and RT-qPCR mixtures contained forward primer (100 to 1000 nM), reverse primer (100 to 1000 nM), and 80 to 400 nM of probe.

Reactions were conducted in triplicate using a Bio-Rad CFX96 thermal cycler (Bio-Rad, California, USA). Each RT-qPCR/qPCR run included triplicate negative and positive controls utilizing gBlock gene fragments (IDT, Coralville, IA, USA). Standards, ranging from 3 × 10^6^ to 3 gene copies (GC) per reaction and prepared from gBlock gene fragments, were used to quantify fecal markers. The baseline and threshold for all qPCR/RT-qPCR assays were consistently set at between 50 and 175 relative fluorescence units (RFU).

### 2.5. Quality assurance/control protocols

The performance characteristics of the qPCR/RT-qPCR assays, including amplification efficiencies (E), correlation coefficient (R²), and y-intercepts, are detailed in Supplementary Table ST4, following the Minimum Information for Publication of Quantitative Real-Time PCR Experiments (MIQE) guidelines (Bustin et al., 2009). The assay limit of detection (ALOD) was determined as the lowest gene copy number with a 95% detection probability for each assay (Verbyla et al., 2016).

A murine hepatitis virus (MHV) RT-PCR assay or a Sketa22 PCR assay was employed to determine PCR inhibition in extracted nucleic acid samples by seeding a known copy number (10^4^) of MHV RNA or *Oncorhynchus keta* (*O. keta*) DNA (Besselsen et al., 2002; Haugland et al., 2005). The reference quantification cycle (Cq) value was determined from triplicate RT-PCR/PCR reactions containing only the positive control and compared with the Cq values obtained from all nucleic acid samples. Samples were considered to have no PCR inhibition when the Cq values of nucleic acid samples were within 2 Cq values of the reference Cq value (Ahmed et al., 2020). Inhibited nucleic acid samples were diluted fivefold or tenfold and then re-analyzed. Nucleic acid extraction and the RT-qPCR/qPCR setup were performed in separate laboratories to minimize contamination introduced in experiments. The membrane blank, filter blank and reagent blank were included for each batch of samples.

#### Data analysis

The accumulation of fecal markers by the passive samplers was characterized using a one compartment, first order kinetic model:

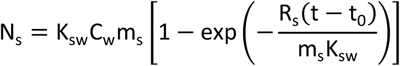

The time for the GC of fecal markers to reach equilibrium (t_eq_) was defined as the time required to attain 99% of the maximal viral load, and was calculated by:

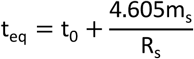

where N_s_ is the log_10_ GC for each target detected from the sampler (log_10_ GC/sampler), K_sw_ is the sampler-water partition coefficient for the fecal markers (log_10_ GC/sampler)/ (log_10_ GC/mL), C_!_ is the concentration of fecal markers in water samples (log_10_ GC/mL), m_s_ is the number of samplers considered in the accumulation model (m = 1), R_s_ is the sampling rate of a passive sampler (mL/h), t is the total deployment duration (h) and t_0_ is the last time point where no quantifiable viral load was observed. Values of R_s_ and K_sw_ were derived from a nonlinear least-squares regression using Graphpad Prism Version 10.4.2 (GraphPad Software, LaJolla, LA).

The root-mean-square error (RMSE), R^2^ and the 95% confidence intervals were used to estimate the goodness of fit.

## 3. Results

### 3.1. qPCR and RT-qPCR assay performance characteristics

The correlation coefficients (R^2^) of RT-qPCR and qPCR standard curves (1 × 10^6^ to 1 GC/μL) for *Carjivirus*, PMMoV, ToBRFV, and GFD ranged between 0.98 and 0.99 (Supplementary Table ST3). The Y-intercepts of standard curves ranged from -42.1 to -37.9. The RT-qPCR and qPCR efficiencies were 101 % for *Carjivirus*, 99.1% for PMMoV, 94.2 % for ToBRFV, and 99.3 % for GFD. The ALODs of *Carjivirus*, PMMoV, ToBRFV, and GFD were 2.4, 6.8, 5.9, and 1.1 GC/reaction, respectively (Table ST4). PCR inhibition was only observed for one grab sample at 0 h and the 72-h composite sample. Blanks and PCR negative control samples did not produce any amplification and were classified as no detection.

### 3.2. Concentrations of host-associated fecal markers in river water samples

*Carjivirus* and GFD were consistently detected in all grab samples collected throughout the study. PMMoV and ToBRFV were detected in all grab water samples except for the sample collected at 0 h. The mean concentrations of *Carjivirus*, PMMoV, ToBRFV, and GFD were 3.96 ± 0.18 log_10_ GC/100 mL, 2.09 ± 0.86 log_10_ GC/100 mL, 2.94 ± 1.22 log_10_ GC/100 mL, and 2.34 ± 0.18 log_10_ GC/100 mL, respectively (Table 1).

**Table 1.** Concentrations of fecal markers i<u>n</u> river water samples.

| Fecal markers | Target concentrations using different sampling methods (mean ± SD) | | |
| --- | --- | --- | --- |
|  | Grab samples (log<sub>10</sub> GC/100 mL) | 24-h time-weighted samples (log<sub>10</sub> GC/100 mL)<sup>a</sup> | 72-h composite sample (log<sub>10</sub> GC/100 mL)<sup>b</sup> |
| *Carjivirus* | 3.96 ± 0.18 | 3.77 ± 0.02 | 4.06 |
| PMMoV | 2.09 ± 0.86 | 2.85 ± 0.02 | 2.66 |
| ToBRFV | 2.94 ± 1.22 | 3.87 ± 0.21 | 3.69 |
| GFD | 2.34 ± 0.18 | 2.34 ± 0.08 | 2.58 |

<sup>a</sup> Mean concentrations and standard deviations were calculated using 24 h time weighed samples collected from 0 - 24 h and 24 - 48 h; <sup>b</sup> Data is based on one 72 h composite sample.

This was the first time the Sipper autosampler, a newly developed submersible autosampler, was used to quantify levels of fecal markers in a river system. A key advantage of the Sipper is that being submersible it can be deployed discreetly underwater, reducing visibility, and minimising the risk of vandalism, a common issue with larger, more exposed autosamplers. Concentrations of two 24-h time-weighted Sipper water samples (0 to 24 h and 24 to 48 h) were consistent for all four markers, indicating little fluctuation in their concentrations over two consecutive days. Only a slight difference was observed between the mean concentrations of the two 24-h time-weighted Sipper water samples collected and the 72-h composite water sample for the four fecal markers, with concentrations expressed as GC/100 mL within a factor of two of each other (Table 1). However, data from the final 24 h time-weighted Sipper sample (48 to 72 h) was not considered since although it was still operating on sample retrieval, the target volume sampled was not achieved, suggesting it was likely collecting less volume per unit time than intended. Due to this uncertainty, the mean of all grab samples was used for calculations involving concentrations in water samples.

### 3.3. Accumulation of fecal markers in passive samplers

All fecal markers exhibited similar accumulation patterns in passive samplers over the 72-h deployment period (Fig. 1). This pattern comprised a relatively short lag time during which marker levels were below the ALOD, followed by increasing accumulation until equilibrium is attained, characterised by equal rates of viral adsorption and desorption to and from sampler membranes.

**Figure 1.**
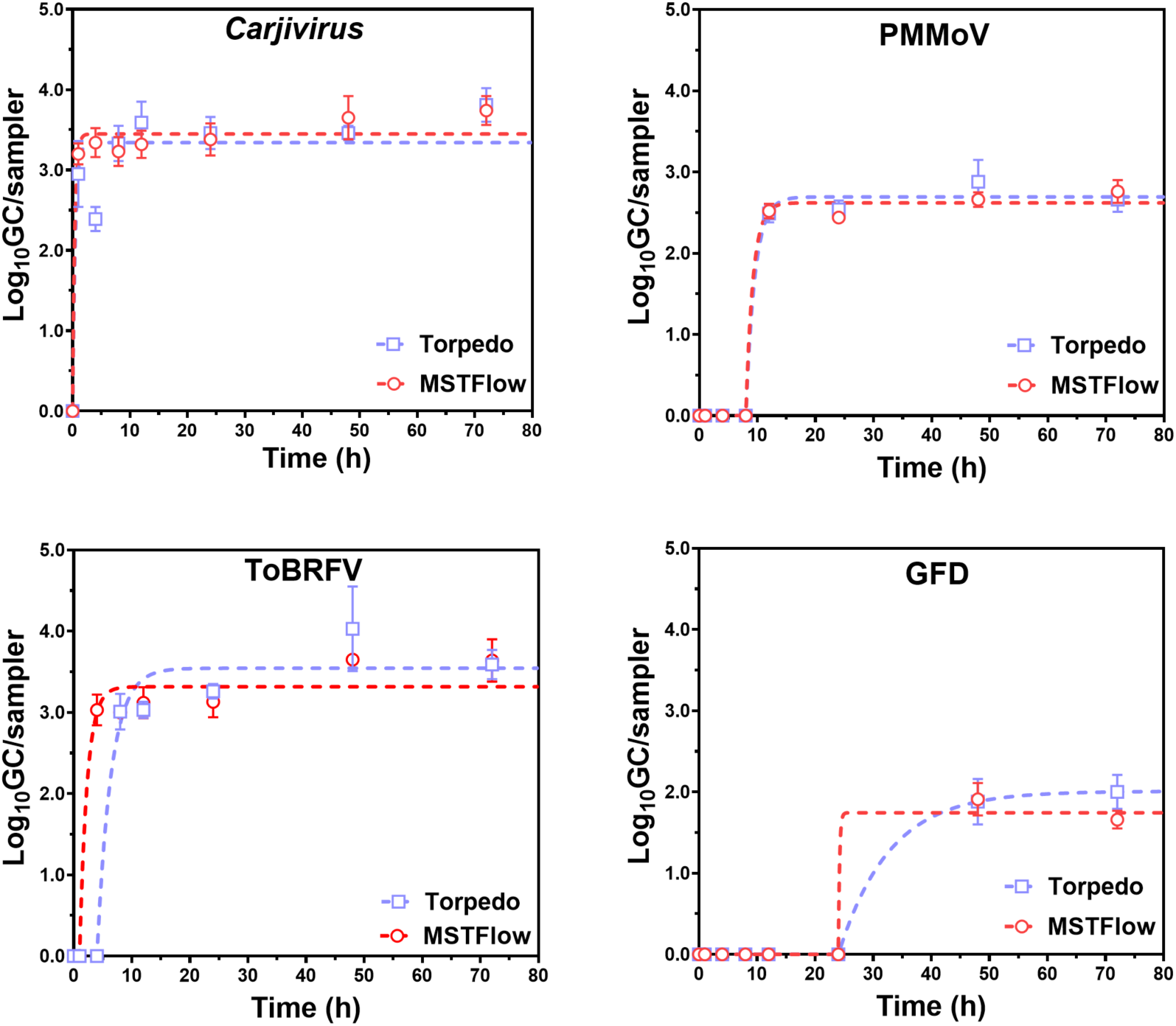
Passive sampler (Torpedo and MSTFlow) fecal marker accumulation profile for riverine deployment.

For PMMoV, a lag time (between 8 and 12 h) was observed in both types of samplers. After this time, viral loads increased with effective equilibrium being attained in less than 16 h. (Table 2). The model-derived sampler/water equilibrium constants (K_“!_) were 7.71 and 7.92 (log_10_ GC/sampler)/ (log_10_ GC/mL) in MSTFlow and Torpedo samplers, respectively. Given mean aqueous levels of 0.34 log_10_ GC/mL, maximal concentrations of PMMoV (K_sw_ x C_w_) were 2.62 and 2.69 log_10_ GC/sampler, respectively. To better appreciate the accumulation capacity of the samplers, using the MSTFlow data as an example, when expressed on a non-logarithmic basis, C_!_= 10^0.34^ = 2.19 GC/mL and the maximal viral load N_“_/*m*_#_ = 10^2.62^ = 416.9 GC/sampler meaning the sampler accumulates over 190 times the amount of virus in each mL of water. An analogous treatment for the Torpedo sampler produces a similar result.

**Table 2.** Detection frequency and concentration of host-associated fecal markers in passive samples collected using Torpedo and MSTFlow samplers with time.

| Target | Sampling time | Torpedo |  | MSTFlow |  |
| --- | --- | --- | --- | --- | --- |
|  |  | Positives (%) | <sup>b</sup>log<sub>10</sub> GC (mean ± SD) | Positives (%) | <sup>b</sup>log<sub>10</sub> GC (mean ± SD) |
| *Carjivirus* | 1 h | 66.7 (6/9) | 2.95 ± 0.41 | 100 (9/9) | 3.20 ± 0.13 |
|  | 4 h | 66.7 (6/9) | 2.39 ± 0.15 | 100 (9/9) | 3.34 ± 0.18 |
|  | 8 h | 100 (9/9) | 3.33 ± 0.22 | 100 (9/9) | 3.23 ± 0.18 |
|  | 12 h | 100 (9/9) | 3.59 ± 0.26 | 100 (9/9) | 3.32 ± 0.17 |
|  | 24 h | 100 (9/9) | 3.46 ± 0.20 | 100 (9/9) | 3.38 ± 0.20 |
|  | 48 h | 100 (9/9) | 3.46 ± 0.09 | 100 (9/9) | 3.66 ± 0.27 |
|  | 72 h | 100 (9/9) | 3.81 ± 0.21 | 100 (9/9) | 3.74 ± 0.18 |
| PMMoV | 1 h | 77.8 (7/9) | DNQ | 88.9 (8/9)<sup>a</sup> | DNQ |
|  | 4 h | 66.7 (6/9) | DNQ | 100 (9/9) | DNQ |
|  | 8 h | 100 (9/9) | DNQ | 100 (9/9) | DNQ |
|  | 12 h | 100 (9/9) | 2.49 ± 0.11 | 100 (9/9) | 2.52 ± 0.09 |
|  | 24 h | 100 (9/9) | 2.54 ± 0.11 | 100 (9/9) | 2.44 ± 0.03 |
|  | 48 h | 100 (9/9) | 2.88 ± 0.27 | 100 (9/9) | 2.66 ± 0.09 |
|  | 72 h | 100 (9/9) | 2.66 ± 0.15 | 100 (9/9) | 2.76 ± 0.14 |
| ToBRFV | 1 h | 22.2 (2/9) | DNQ | 44.4 (4/9) | DNQ |
|  | 4 h | 11.1 (1/9) | DNQ | 66.7 (6/9) | 3.03 ± 0.19 |
|  | 8 h | 100 (9/9) | 3.01 ± 0.22 | 100 (9/9) | 3.00 ± 0.05 |
|  | 12 h | 100 (9/9) | 3.03 ± 0.10 | 100 (9/9) | 3.12 ± 0.19 |
|  | 24 h | 100 (9/9) | 3.26 ± 0.09 | 88.9 (8/9) | 3.13 ± 0.19 |
|  | 48 h | 100 (9/9) | 4.03 ± 0.52 | 100 (9/9) | 3.65 ± 0.06 |
|  | 72 h | 100 (9/9) | 3.59 ± 0.18 | 100 (9/9) | 3.64 ± 0.26 |
| GFD | 1 h | 0 (0/9) | DNQ | 0 (0/9) | DNQ |
|  | 4 h | 11.1 (1/9) | DNQ | 11.1 (1/9) | DNQ |
|  | 8 h | 11.1 (1/9) | DNQ | 11.1 (1/9) | DNQ |
|  | 12 h | 11.1 (1/9) | DNQ | 22.2 (2/9) | DNQ |
|  | 24 h | 44.4 (4/9) | DNQ | 44.4 (4/9) | DNQ |
|  | 48 h | 77.8 (7/9) | 1.88 ± 0.28 | 66.7 (6/9) | 1.91 ± 0.20 |
|  | 72 h | 66.7 (6/9) | 2.00 ± 0.21 | 77.8 (7/9) | 1.66 ± 0.11 |

<sup>a</sup>Replicates include three biological replicate samplers, each contains three PCR technical replicates; <sup>b</sup>Mean GC/membrane was calculated when at least two out of three biological replicate samples were quantifiable; DNQ: did not quantify.

For ToBRFV, lag times were 1 h and 4 h in MSTFlow and Torpedo samplers respectively, with effective equilibrium attained after a further 5.9 and 11.2 h. Sampler/water equilibrium constants (K_“!_= 2.79 and 2.98 (log_10_ GC/sampler)/ (log_10_ GC/mL)) were decreased compared to those observed with PMMoV due to river levels of this viral fecal marker being much larger (1.19 log_10_ GC/mL). At equilibrium when maximal loads were attained, concentrations were 3.32 and 3.54 log_10_ GC/sampler in MSTFlow and Torpedo samplers, respectively. Consequently, when expressed on a non-logarithmic basis, the MSTFlow sampler accumulates over 130 times the amount of virus in each mL at equilibrium and the Torpedo sampler over 220 times.

*Carjivirus* was quantifiable in all qPCR replicates for both types of passive sampler, but no lag time was observed. Sampler/water equilibrium constants (K_“!_= 1.76 and 1.71 (log_10_ GC/sampler)/ (log_10_ GC/mL) in MSTFlow and Torpedo samplers were reduced compared to those of the other human fecal markers while mean aqueous concentrations were the highest (1.96 log_10_ GC/mL). Equilibrium viral loads were 3.45 and 3.35 log_10_ GC/sampler in MSTFlow and Torpedo samplers, respectively. Consequently, when expressed on a non-logarithmic basis, sampler equilibrium concentrations were 30 and 23 times that in one mL of river water. Attainment of equilibrium was however the most rapid of all viruses of interest, being < 2.5 h with both types of samplers.

In contrast, the accumulation of the bacterium GFD on passive sampler was comparatively slow, with no detection observed in either sampler type during the first 24 h of deployment. Effective equilibrium (2.01 log_10_ GC /sampler) for the Torpedo sampler was not attained until 64 h after initial deployment. Lack of data prevented the estimation all model parameters for the MSTFlow sampler. At equilibrium, concentrations in the sampler (10^2.01^ = 102.3 GC/sampler) were over 40 times that in a mL of river water (10^0.34^ = 2.19 GC/mL).

#### Sampling Kinetics for the MSTFlow and Torpedo passive samplers

Based on the accumulation curves, the uptake of *Carjivirus*, PMMoV, and ToBRFV onto passive membranes over 72 h was fitted using a first order one compartment kinetic model with a prior lag time. For GFD, only data for the Torpedo sampler produced stable fitted parameters when the model was applied. The estimated sampling rates (R_s_) and effective times to equilibrium for *Carjivirus*, PMMoV, and ToBRFV are presented in Table 3. The mean of the individual grab samples was used to represent the average time-weighted concentration during deployment and subsequently for deriving sampling kinetics. The sampling rates (mL/h) were lowest for GFD (0.68 for the Torpedo sampler) compared to those for the viral fecal markers (6.29 and 5.11, 2.17 and 1.22, 4.63 and 3.63 for MSTFlow and Torpedo samplers respectively with PMMoV, ToBRFV and *Carjivirus*).

**Table 3.** Estimated sampling rates (R_s_, mL/h), times to equilibrium conditions (t_eq_, h), and sampler-water partition coefficients (K_sw_) for Torpedo and MSTFlow passive samplers based on a first-order kinetic model.

| Parameters | *Carjivirus* |  | PMMoV |  | ToBRFV |  | GFD |
| --- | --- | --- | --- | --- | --- | --- | --- |
|  | Torpedo | MSTFlow | Torpedo | MSTFlow | Torpedo | MSTFlow | Torpedo |
| $R_s$ (mL/h) | 3.63 | 4.63 | 5.11 | 6.29 | 1.22 | 2.17 | 0.68 |
| $t_{eq}$ (h) | 2.2 | 1.7 | 15.1 | 13.6 | 15.2 | 6.9 | 64.2 |
| $K_{sw}$ ($\log_{10}$ GC/sampler)/($\log_{10}$ GC/mL) | 1.71 | 1.76 | 7.92 | 7.71 | 2.98 | 2.79 | 5.91 |
| $^a R^2$ | 0.46 | 0.76 | 0.99 | 0.99 | 0.95 | 0.95 | 0.98 |
| RMSE | 0.48 | 0.25 | 0.15 | 0.11 | 0.34 | 0.28 | 0.12 |
| $^b 95\%$ CI $R_s$ | 2.57 to 7.17 | 3.70 to 6.87 | 4.26 to 6.78 | 5.15 to 9.99 | 0.99 to 1.63 | 1.70 to 3.26 | 0.52 to 1.67 |
| $^b 95\%$ CI $K_{sw}$ | 1.64 to 1.77 | 1.72 to 1.79 | 7.75 to 8.10 | 7.58 to 7.83 | 2.87 to 3.09 | 2.72 to 2.86 | 5.64 to 6.21 |

<sup>a</sup> The $R^2$ value indicates the goodness of fit for each model, where higher $R^2$ values indicates a better fit. <sup>b</sup> 95% CI $R_s$ and $K_{sw}$ are the 95% confidence intervals.

Derived sampling rates (R_s_) and sampler/water partition coefficients (K_sw_) for MSTFlow and Torpedo samplers were not significantly different with both PMMoV (F(2,124) = 2.10, *P* = 0.13) and *Carjivirus* (F(2,122) = 1.66, *P* = 0.19). Comparable assessments were unable to be undertaken for ToBRFV and GFD because of differing lag times and lack of data respectively. These results suggest a close correspondence between times to equilibrium for samplers with PMMoV and *Carjivirus* (Table 3), which is expected since sampling rates are a factor in deriving the effective time to equilibrium along with K_sw_ values. Consequently, the fact that there is a marked difference in equilibrium times for ToBRFV (15.2 h and 6.9 h for Torpedo and MSTFlow samplers) reflects contributing R_“_ values or K_sw_ values or both being different for passive samplers with this fecal marker.

## 4. Discussion

In this study, three human fecal markers (*Carjivirus*, PMMoV, and ToBRFV) and one avian fecal marker (GFD) were consistently present in water samples throughout the sampling period. The prevalence of human fecal markers observed in water samples was not unexpected as the samples were collected downstream of a sewage infrastructure pipeline, and a 24 h precipitation event had occurred prior to sampling that might have contributed the release of untreated wastewater from the pipeline (i.e., high rainfall for 3 d finishing 1 d before the sampling, Table S1). The *Helicobacter* spp. associated GFD is highly specific to avian host groups and has been validated as a reliable marker to detect avian fecal pollution in environmental waters (Ahmed et al., 2016). The consistent detection of GFD suggested ongoing bird fecal pollution in the study area, with water birds likely being a major source of pollution.

Viral adsorption (or their DNA/RNA fragments) to a solid surface can be conceptualized as a two-step process in which the virions/fragments are first transported close to the adsorbent surface and then immobilized there by electrostatic, dispersive, and/or possibly chemical interactions (Gerba, 1984; Bitton, 1975). These two steps largely determine the overall adsorption rate, which relies on multiple parameters, including the concentration of viruses in direct contact with the adsorptive surface, the diffusion velocity of virions, the interfacial surface area, and the availability of adsorption sites (Grand et al., 1993). In passive sampling, Schang et al. (2021) found a statistically significant correlation between the daily mean log_10_ GC concentration of SARS-CoV-2 measured in wastewater and the mean log_10_ GC of SARS-CoV-2 load on the passive samplers, regardless of the sampling materials (Schang et al., 2021). In a study on bacteriophage MS2 in deionized water and wastewater using negatively charged membranes, higher initial concentrations were associated with increased adsorption (Shakallis et al., 2024). In the current work, the relatively higher accumulated concentrations of *Carjivirus* and ToBRFV on passive sampler corresponded with higher concentrations in grab water samples. Although a correlation analysis was not performed due to insufficient data, this pattern supports the expected accumulation behavior of passive samplers, which can be interpreted within the context of the kinetic model.

A first order one compartment kinetic model with a prior lag time was employed to describe the adsorption of the viral fecal markers *Carjivirus*, PMMoV, and ToBRFV, providing a quantitative approach to understanding their uptake dynamics in passive samplers. This study observed distinct sampling behaviors between wastewater and river water, with a notable lag phase in viral (PMMoV and ToBRFV) uptake from the river that was not previously reported during previous wastewater deployments (Li et al., 2022). In river water, the accumulation transitioned to operation in equilibrium mode within 16 h of initial deployment including lag times. In contrast, viral accumulation in wastewater exhibited a linear profile beyond 48 h despite higher viral concentrations.

The lag phase in river water may arise from electrostatic interactions between viral particles and the sampler’s mixed cellulose ester membrane (cellulose nitrate and acetate). This membrane is historically described as “negatively charged” due to residual carboxylate groups (COO⁻). In river water (pH 7.4) (Supplementary Table ST2), viruses with isoelectric points (pI) of 3 to 6 carry a negative charge, potentially causing coulombic repulsion with the membrane (Michen and Graule. 2010). PMMoV has a reported pI of 3.3 to 3.8 (Dhakar and Geetanjali. 2022). Although the pI of ToBRFV has not been reported, its structural similarity to other members of the *Tobamovirus* genus-such as Tobacco mosaic virus (TMV) which has a reported pI of approximately 3.5 suggests that ToBRFV likely possesses a similarly acidic pI (Oster, 1951). This repulsion could delay initial sorption until alternative mechanisms such as hydrophobic interactions or physical entrapment facilitate accumulation, initiating quantifiable uptake after the lag phase. In wastewater, the observed linear viral uptake may be due to the high organic matter content reducing or mitigating electrostatic barriers. Organic matter may coat the membrane or viral particles, reducing repulsion by shielding charged groups or altering surface properties. However, wastewater’s complex matrix, rich in dissolved organic carbon, suspended solids, and surfactants, likely also slows mass transfer through fouling or competition for binding sites, extending the linear uptake phase.

*Carjivirus* with a pI of 3.8, exhibited a relatively rapid adsorption on the membranes with effective equilibrium attained within 2.5 h, and no lag phase was observed. *Carjivirus* is characterized by an icosahedral capsid structure with a short, hydrophobic tail, similar to other tailed bacteriophages which likely contributes to its comparatively high surface attachment efficiency (Heffron and Mayer., 2021). Additionally, the relatively high concentration of *Carjivirus* in river waters may also facilitate its adsorption, as observed in another study (Hayes et al., 2022).

The avian bacterial marker GFD was comparatively less abundant in river water samples, exhibiting a prolonged lag phase (at least 24 h) with the lowest accumulated concentrations on passive sampler membranes. *Helicobacter* spp. are gram-negative, curved to spiral-shaped bacteria characterized by a net negative surface charge, primarily attributed to the presence of lipopolysaccharides in their outer membrane (O’Toole and Clyne., 2001). As bacteria (e.g., *Helicobacter* spp.) generally possess a larger size relative to viruses, it would be expected that the larger size results in a lower aqueous diffusion coefficient and, consequently, a longer time required to encounter surfaces for potential adsorption (Miller., 1924). Additionally, the larger size of bacteria may subject them to greater hydrodynamic drag in flowing systems, thereby limiting their ability to approach surfaces closely enough for effective adsorption, in contrast to smaller, more agile viruses (Margalit et al., 2013). Given that GFD was only detected at the 48 h and 72 h time points in the passive sampler deployment, an extended deployment duration may be necessary to facilitate its adsorption and equilibrium on the negatively charged membrane.

The sampling rates for the two of the three viral targets (*Carjivirus* and PMMoV), defined by the volume of water from which the analyte is extracted per unit of time (mL/h), were not significantly different between MSTFlow and Torpedo passive samplers. The difference in K_sw_ values between the two samplers for both these viruses was also negligible, which may be expected since the sampling materials (adsorbent membranes) were the same for both, and environmental conditions remained consistent for both samplers.

The housing for both devices were primarily designed to protect the enclosed membrane filters which are fragile and susceptible to physical degradation under high flow conditions but allow sufficient water flow while also preventing high ragging rates (the accumulation of many solid particles from water) that could limit interactions between the passive sampler and target microbes (Schang et al., 2021; Hayes et al., 2021). In this study, no particles were observed on the passive sampler membranes following deployment (river water turbidity was 61.6 NTU). Therefore, the effect of particles (e.g., total suspended solids) could be considered negligible, while the 72-h deployment duration likely allowed for sufficient interactions and equilibrium attainment between the passive membranes and target viral fecal markers.

This study provides first-hand data on the in-situ performance and calibration of membrane-based passive sampling for fecal markers for environmental water quality monitoring. Furthermore, passive sampling could effectively reduce the number of samples that need to be collected and processed when in-field calibration data are available, enabling large-scale passive sampling deployment in the future. Passive sampling also demonstrated relatively high sensitivity for detecting pathogens present at low abundances. Especially for PMMoV and ToBRFV, which had concentration factors greater than 200, a passive sampler detecting 50 GC per membrane would correspond to a grab sample requiring more than 1 liter of water assuming no losses during sample concentration and extraction to detect an equivalent amount (see Supplementary Table ST5).

While the study yielded promising results, several challenges remain. For instance, it focused on a relatively urbanized river environment, and the effectiveness of passive samplers in more complex or high turbidity settings requires further exploration. In their current configuration, the samplers reach equilibrium within hours, reflecting concentrations over a relatively short time. This equilibrium mode of sampling may limit their ability to capture temporal fluctuations in environments with highly variable concentrations, unlike kinetic mode sampling, which is better suited for such conditions.

The first-order kinetic model used to characterize fecal marker accumulation relies on several assumptions that may affect its accuracy. First, it assumes constant sampler-water partition coefficient (Ksw) and sampling rate (Rs), which may vary due to environmental factors such as flow, turbidity or biofouling. Second, the model assumes stable water concentrations (Cw), but precipitation suggests temporal variability from sewage inputs or dilution, potentially impacting accumulation, especially for PMMoV and ToBRFV with observed lag phases. Third, it assumes no desorption of markers, which may not hold for *Carjivirus* or GFD, possibly underestimating (Ns). Fourth, the single-sampler assumption (ms = 1) may overlook design-specific effects, as seen with ToBRFV’s differing lag times. Finally, log_10_ transformations assume linear behavior in log space, but lower R² for *Carjivirus* (0.46 for Torpedo and 0.76 for MSTFlow) suggests potential non-linearities. These limitations highlight the need for cautious interpretation and future validation through experiments assessing biofouling, variable (Cw), or alternative models to enhance passive sampling reliability.

Future research should focus on extending the duration of kinetic mode to enable prolonged time-integrative sampling for more comprehensive monitoring in dynamic environments. Moving forward, a key priority will be developing a passive sampling framework that can be widely applied in large-scale environmental monitoring programs. Future research should also compare the performance of passive samplers with different adsorption materials across various water matrices.

## 5. Conclusions

- The findings of this study highlight that membrane-based passive samplers are effective tools for detecting microbial contaminants in urban river environments.
- Passive sampling methods not only offer a more efficient, sensitive, cost-effective, and complementary technique to traditional water quality monitoring techniques, but also present a promising tool for advancing environmental pathogen surveillance and safeguarding public health.
- The study contributes to the growing body of literature the supports passive sampling and sets the stage for further research into its application across diverse aquatic environmental systems and microbial targets.

