## Supplementary Tables and Figure for "In-situ calibration of passive samplers for monitoring host-associated fecal markers in an urban river"

[1]CSIRO Environment, Ecosciences Precinct, 41 Boggo Road, Dutton Park, QLD 4102, Australia.

[2]Queensland Alliance for Environmental Health Sciences (QAEHS), The University of Queensland, 20 Cornwall Street, Woolloongabba, QLD 4103, Australia.

[3]School of Environment and Science, Griffith University, Nathan 4111 Queensland, Australia.

**Supplementary Table ST1** Precipitation data from November 2024 to December 2024 in sampling area.

| Date | Precipitation (mm) | |
|---|---|---|
| | November | December |
| 1st | 0 | 20.2 |
| 2nd | 9.0 | 55.2 |
| 3rd | 0.8 | 2.2 |
| 4th | 0.2 | 1.6 |
| 5th | 0.6 | 14.2 |
| 6th | 3.2 | 0 |
| 7th | 0 | 0 |
| 8th | 0 | 0 |
| 9th | 0 | 0.2 |
| 10th | 0 | 0 |
| 11th | 17.4 | 75.8 |
| 12th | 10.0 | 0.2 |
| 13th | 0 | 0 |
| 14th | 28.4 | 1.2 |
| 15th | 1.0 | 70.2 |
| 16th | 0.2 | 0.8 |
| 17th | 68.6 | 59.4 |
| 18th | 0 | 5.6 |
| 19th | 2.6 | 13.8 |
| 20th | 37.4 | 0 |
| 21st | 7.4 | 0 |
| 22nd | 8.4 | 0 |
| 23rd | 24.0 | 0 |
| 24th | 0.6 | 0 |
| 25th | 0 | 0 |
| 26th | 0 | 0 |
| 27th | 2.0 | 0 |
| 28th | 1.0 | 0 |
| 29th | 0.4 | 16.8 |
| 30th | 23.0 | 0 |
| 31st | | 1.6 |

**Supplementary Table ST2** Water physico-chemical parameters in the sampling area.

| Parameters (unit) | Value |
|---|---|
| Temperature (℃) | 26.6 |
| Conductivity (25 ℃ mS/cm) | 2.2 |
| Salinity (g/L) | 1.1 |
| pH | 7.4 |
| DO Sat (%) | 52.7 |
| DO (mg/L) | 4.2 |
| Secchi depth (m) | 0.3 |
| Turbidity (NTU) | 61.6 |
| Chlorophyll-a (μg/L) | 4.9 |
| Phaeopigments (μg/L) | 0.1 |
| Total nitrogen (mg/L) | 1.3 |
| Nitrogen Ammonia (mg/L) | 0 |
| Nitrogen Oxidised (mg/L) | 0.4 |
| Phosphorus Total (mg/L) | 0.3 |
| Phosphorus Filterable Reactive (mg/L) | 0.2 |
| TSS (mg/L) | 55 |

**Supplementary Table ST3** RT-qPCR and qPCR primer and probe sequences, concentrations and cycling parameters used in this study.

| Target | Oligonucleotides sequences (5' - 3') | Primers and probes (nM) | Cycling parameters | References |
|---|---|---|---|---|
| *Oncorhynchus keta* | F: GGT TTC CGC AGC TGG G | 500 | 95 °C for 10 min and 45 cycles of 95 °C for 15 s, 63 °C for 45 s | Haugland et al., 2005 |
| | R: CCG AGC CGT CCT GGT CTA | 500 | | |
| | P: FAM-AGT CGC AGG CGG CCA CCG T-TAMRA | 400 | | |
| Murine hepatitis virus (MHV) | F: GGA ACT TCT CGT TGG GCA TTA TAC T | 300 | 50°C for 10 min for RT; 95°C for 5 min and 45 cycles of 95°C for 15 s, 60°C for 60 s | Besselsen et al., 2002 |
| | R: ACC ACA AGA TTA TCA TTT TCA CAA CAT A | 300 | | |
| | P: FAM-ACA TGC TAC GGC TCG TGT AAC CGA ACT GT-BHQ | 400 | | |
| *Carjivirus* | F: CAG AAG TAC AAA CTC CTA AAA AAC GTA GAG | 1000 | 95°C for 10 min and 45 cycles of 95°C for 15 s, 60°C for 60 s. | Stachler et al., 2017 |
| | R: AT GAC CAA TAA ACA AGC CAT TAG C | 1000 | | |
| | P: FAM-AAT AAC GAT TTA CGT GAT GTA AC-TAMRA | 100 | | |
| Pepper mild mottle virus (PMMoV) | F: GAG TGG TTT GAC CTT AAC GTT TGA | 200 | 50°C for 10 min for RT; 95°C for 10 min and 45 cycles of 95°C for 30 s, 53°C for 60 s and 72°C for 60 s | Rosario et al., 2009; Haramoto et al., 2013 |
| | R: TTG TCG GTT GCA ATG CAA GT | 200 | | |
| | P: FAM-CCT ACC GAA GCA AAT G-MGBNFQ | 80 | | |
| Tomato Brown Rugose Fruit Virus (ToBRFV) | F: TCA GTG TCT GTT TGG TCG ATA A | 500 | 50°C for 10 min for RT; 95°C for 10 min and 45 cycles of 95°C for 30 s, 57.7°C for 30 s | Natarajan et al., 2023 |
| | R: GGA ACG ACT TTG AAC TGA AAC C | 500 | | |
| | P: FAM-AGA GCG GAC GAG GCA ACT CTT G | 500 | | |
| *Helicobacter* spp. GFD | F: TCG GCT GAG CAC TCT AGG G | 100 | 10 min at 95°C, 45 cycles of 10 s at 95°C, 30 s at 57°C, 20 s at 72°C | Green et al., 2012 |
| | R: GCG TCT CTT TGT ACA TCC CA | 100 | | |
| | P: FAM-AAG GAG GAG GAA GGT GAG GAC GA-BHQ1 | 100 | | |

**Supplementary Table ST4** qPCR and RT-qPCR performance characteristics.

| Assays | Performance characteristics | | | | |
|---|---|---|---|---|---|
| | Efficiency (E) (%) | Linearity ($R^2$) | Slope | Y-intercept | ALOD (GC/reaction) |
| *Carjivirus* | 101 | 0.99 | -3.31 | 40.1 | 2.4 |
| PMMoV | 99.1 | 0.99 | -3.34 | 37.4 | 6.8 |
| ToBRFV | 94.2 | 0.99 | -3.47 | 42.0 | 5.9 |
| GFD | 99.3 | 0.98 | -3.34 | 37.9 | 1.1 |

**Supplementary Table ST5** Illustration of passive samplers' sensitivity compared to grab water samples.

| Parameters | *Carjivirus* | PMMoV | ToBRFV | GFD |
|---|---|---|---|---|
| Concentration factor | 23 - 30 | 190 - 230 | 130 - 220 | 40 |
| [a]$C_p$ (GC/sampler) | 50 | 50 | 50 | 50 |
| [b]Cw (GC/mL) in grab sample | 1.67 - 2.17 | 0.22 - 0.26 | 0.23 - 0.38 | 1.25 |
| ALOD (GC/reaction) | 2.4 | 6.8 | 5.9 | 1.1 |
| [c]Extracted/eluted volume in tube (µL) | 150 | 150 | 150 | 150 |
| [d]Reaction volume for qPCR (µL) | 3 | 3 | 3 | 3 |
| [e]$V_w$ Least sampling volume for grab sample (mL) | 56 - 72 | 1292 - 1564 | 767 - 1298 | 44 |

[a]Assumed concentrations in passive sampler for each target; [b]Concentrations in grab water sample were derived using the assumed concentrations in passive sampler divided by the concentration factor; [c]Extracted/eluted volume is adopted from methods used in the paper, please see Section 2.3; [d]Reaction volume for qPCR is adopted from methods used in the paper, please see Section 2.4; [e]Value is calculated assuming no loss occurred during sample concentration and extraction, the equation is: $V_w = \frac{(ALOD/3)}{150} \div C_w$.

(a)

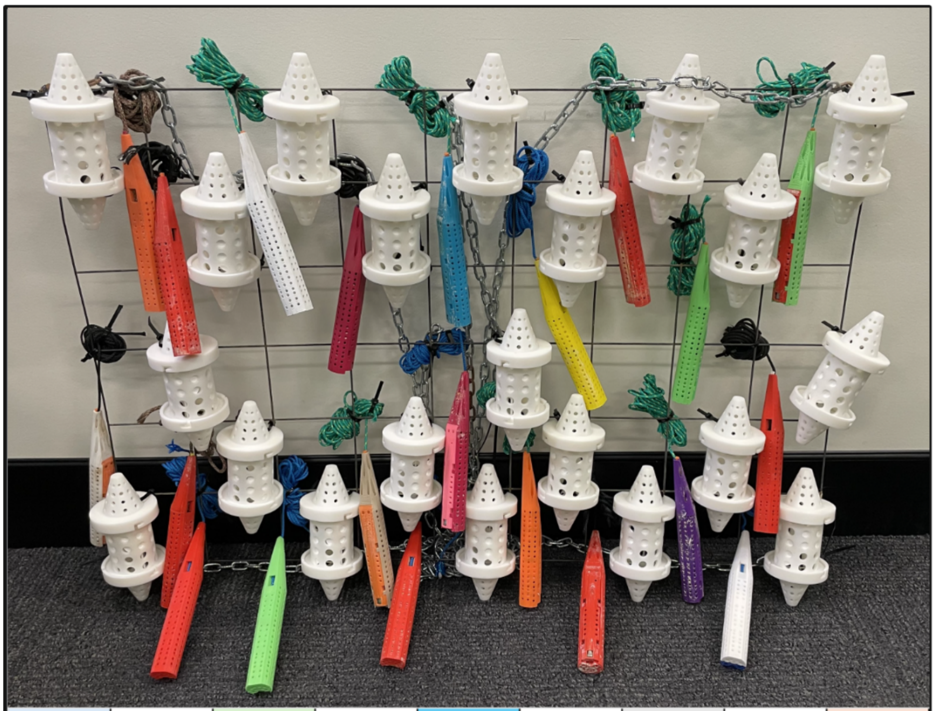

(b)

| | | | | | | | | |
|---|---|---|---|---|---|---|---|---|
| 4 h replicate A | | 12 h replicate A | | 72 h replicate B | | 24 h replicate A | | 8 h replicate A |
| | 72 h replicate A | | 4 h replicate B | | 8 h replicate B | | 48 h replicate A | |
| | 1 h replicate A | | | 1 h replicate B | | | | 1 h replicate C |
| | 24 h replicate C | | 48 h replicate B | | 12 h replicate B | | 4 h replicate C | |
| 48 h replicate C | | 8 h replicate C | | 24 h replicate B | | 72 h replicate C | | 12 h replicate C |

**Supplementary Figure SF1** Deployment setup and sampling design of passive samplers (a) Arrangement of passive samplers (Torpedo and MSTFlow) prepared for deployment. (b) Sampling schedule and randomized positioning layout for different exposure times and replicates.
